# The N300: An Index For Predictive Coding Of Complex Visual Objects and Scenes

**DOI:** 10.1101/2020.09.21.304378

**Authors:** Manoj Kumar, Kara D. Federmeier, Diane M. Beck

## Abstract

Predictive coding models can simulate known perceptual or neuronal phenomena, but there have been fewer attempts to identify a reliable neural signature of predictive coding for complex stimuli. In a pair of studies, we test whether the N300 component of the event-related potential, occurring 250-350 ms post-stimulus-onset, has the response properties expected for such a signature of perceptual hypothesis testing at the level of whole objects and scenes. We show that N300 amplitudes are smaller to representative (“good exemplars”) compared to less representative (“bad exemplars”) items from natural scene categories. Integrating these results with patterns observed for objects, we establish that, across a variety of visual stimuli, the N300 is responsive to statistical regularity, or the degree to which the input is “expected” (either explicitly or implicitly) based on prior knowledge, with statistically regular images evoking a reduced response. Moreover, we show that the measure exhibits context-dependency; that is, we find the N300 sensitivity to category representativeness when stimuli are congruent with, but not when they are incongruent with, a category pre-cue. Thus, we argue that the N300 is the best candidate to date for an index of perceptual hypotheses testing for complex visual objects and scenes.

## Introduction

The stars in the night sky are not arranged in the shape of a great bear and there is no rabbit on the moon; it is our prior knowledge of these shapes that invokes such descriptions. Increasingly, it is clear that perception does not depend on the sensory stimulus alone but is also dynamically influenced by our prior knowledge (Smith and Loschky 2019; Gordon et al. 2017; Caddigan et al. 2017; Lupyan 2017; Vo and Wolfe 2013; Voss et al. 2012; Summerfield et al. 2006). Indeed, many models of perception include some form of perceptual hypothesis testing (PHT), in which perception, a hard inverse problem, is conceived of as a process of generating a hypothesis on the basis of both sensory input and prior knowledge and the current context (Clark 2013; Gregory 1980; Hochberg 1981; Huang and Rao 2011; Rock 1983; Helmholtz 1925). Recently, one class of PHT models has garnered increased interest: predictive coding models (Rao and Ballard 1999; Friston 2005; Spratling 2010), which posit that each area of, for example, visual cortex learns statistical regularities from the world that it then uses, jointly with the input from the preceding area, to make predictions about the stimulus. In particular, the prediction and incoming sensory signal are proposed to undergo an iterative matching process at each stage of the processing hierarchy. Most of these models are hierarchical in nature, with the prediction feeding back on the preceding area. The mismatch (“prediction error”), if any, between the prediction and the incoming sensory signal is then propagated to higher layers in the processing hierarchy, revising the weights of the hypotheses, until the feedback matches the incoming signal and the error is zero (Rao and Ballard 1999; Friston 2005; Lange et al. 2018). These predictive coding models have risen to prominence in recent years, in part because they represent an efficient coding scheme for the complexity of the visual world and, perhaps more importantly, because they posit a role for the abundant feedback connections known to exist between visual areas.

The bulk of support for predictive coding models has come from the models’ ability to simulate known perceptual or neuronal phenomena (reviewed in Spratling 2016). The empirical data used for such models have primarily come from experiments manipulating basic features of simple stimuli, such as variations in grating orientation or color (Kok et al. 2017; Marzecová et al. 2017, 2018; Rungratsameetaweemana et al. 2018; Smout et al. 2019, 2020). However, it should also be possible to find signatures of predictive coding at higher levels of visual analysis. Such a signature would be observed to a variety of types of complex visual stimuli (objects, faces, natural scenes) across most or all viewing conditions. More importantly, it should be responsive to statistical regularity, or the degree to which features in the input are “expected” (either explicitly or implicitly) by the system based on prior knowledge. We learn regularities of object and natural scene features by being exposed to prototypical objects and natural environments over our lifetime. This prior knowledge facilitates our processing when the regularities in the incoming sensory stream meet our expectations (Caddigan et al. 2017). Thus, a good measure of predictive coding would index when stimuli deviate from the regularities we expect to see. In particular, the measured response should increase with increasing irregularity, in keeping with the increased iterations, or inference-based error, proposed to occur when an item does not match the prediction. Importantly, the measure should also show context-dependency, as statistical regularities need to be sensitive to the immediate context in order to be of use to the system.

Using complex visual objects, Schendan and colleagues (Schendan and Kutas 2002, 2003, 2007) have shown that the N300 component of the event-related potential (ERP) can be interpreted as an index of object model selection processes, a framework that fits within PHT (Schendan and Ganis 2012; Schendan 2019). Here we build on these findings, addressing the question of whether the N300 is also sensitive to statistical regularity for complex visual stimuli other than objects -- in particular, for good and bad examples of visual scenes. Moreover, critically, we ask whether the N300 is sensitive to in-the-moment expectations for visual information, as established by, in the present work, verbal cues. Taken together, this kind of evidence would support the characterization of the N300 more broadly as a signature of predictive coding mechanisms, operating in occipitotemporal visual cortex at the scale of whole objects and scenes.

### The N300

The N300 is a negative going component with a frontal scalp distribution that peaks around 300 ms after the onset of a visual stimulus. It has been shown to be sensitive to global perceptual properties of visual input (Mcpherson and Holcomb 1999; Schendan and Kutas 2002, 2003) but not to manipulations limited to low level visual features (e.g., color, or small-scale line segments; Schendan and Kutas 2007) that are known to be processed in early visual cortex. Components that precede the N300 in time have instead been linked to processing of and expectations for such low-level features. For example, a component known as the visual mismatch negativity (vMMN) occurs between 100-160 ms in target-oddball paradigms, where it is larger for the visual oddball stimuli. The vMMN has sometimes been associated with predictive coding (Stefanics et al. 2014; Oxner et al. 2019). However, given its sensitivity to the current experimental context – and, importantly, not to statistical regularities built up over a lifetime – as well as its source location to occipital cortex (Susac et al. 2014; File et al. 2017), the vMMN would be classified as indexing early stage PHT processing. In contrast, the N300 is a “late” visual component, with likely generators in occipitotemporal cortex (Schendan, 2019; Sehatpour et al., 2006). It immediately precedes access to multimodal semantic memory (reflected in the N400, which is observed later in time than the N300 when both are present; Kutas and Federmeier 2011). The N300 is therefore well positioned to capture the iterative, knowledge- and context-sensitive process of visual processing of the global features of stimuli, as proposed by predictive coding models, and thus seems promising as a candidate index of intermediate to late stage PHT processing.

Importantly, as hypothesized by predictive coding models, the amplitude of the N300 increases for less “expected” (i.e., less statistically regular) stimuli. The N300 is larger to pictorial stimuli that lack a global structure as compared to when the global structure of the object is clearly discernible (Schendan and Kutas 2003). The N300 is also sensitive to repetition, with a reduced amplitude for repeated presentations; importantly, however, N300 repetition effects (but not those on earlier components) depend on knowledge, as they are larger when the visual stimulus is meaningful (Voss and Paller 2007; Schendan and Maher 2009; Voss et al. 2010). Similarly, and critically, N300 amplitudes are sensitive to a variety of factors that reflect the degree to which an object fits with prior experience. For example, N300 amplitudes are sensitive to the canonical view of an object; an open umbrella oriented horizontally (non-canonical) elicits a larger N300 amplitude than an open umbrella oriented vertically (Schendan and Kutas 2003; Vo and Wolfe 2013). Amplitude modulations are also linked to factors such as object category membership, presence of category-diagnostic object features, and (rated) match to object knowledge (Gratton et al., 2009; Schendan, 2019; Schendan & Maher, 2009). This pattern of data suggests that the N300 may be a good marker for not only the global structure of an object but the degree to which the input matches learned statistical regularities more generally, with larger N300 amplitudes for stimuli that do not match predictions based on learned regularities and hence require further processing.

Thus far, empirical data have largely linked the N300 to object processing, sometimes in the context of a scene (Mudrik et al. 2010; Vo and Wolfe 2013; Lauer et al. 2020), but still ostensibly elicited by an object. Indeed, Schendan (2019) has specifically linked the N300 to object model selection processes, in which an input is matched to possible known objects. This model selection process includes PHT computations. Here, however, we hypothesize that the N300 may reflect a more general signature of hierarchical inference within higher level visual processing. If so, it should be elicited by other meaningful visual stimuli, such as natural scenes. Scenes differ from individual objects in a few ways. Scenes often contain multiple objects rather than prominent objects that overshadow their backgrounds. Moreover, the spatial layout of the environment is much more critical for understanding a photograph of a scene than a photograph of an object. Finally, it is clear that the human visual system sees objects and scenes as importantly different as they have sub-systems dedicated to processing them (Epstein & Kanwisher, 1998). Thus, if the N300 reflects, not a specific facet of object processing but, more generally, the computations associated with PHT in higher level vision, then it should also be sensitive to statistical regularity and prediction during scene processing.

In fact, scrambled scenes (created by recombining parts of the scene image into a random jigsaw) have been found to elicit larger N300 amplitudes compared to intact and identified scenes (Pietrowsky et al. 1996). Because the scrambled scenes were degraded, however, it is not clear whether these effects simply reflect the disruption to the global structure of the image or a deviation from statistical regularity more generally. Here we use intact scenes that are either highly representative of their category (e.g., good exemplars of that category) or less representative of their category (bad exemplars). Importantly, all the images are good photographs of real world scenes (i.e., they are not degraded); they are statistically regular or irregular by virtue of how representative they are of their category. A highly representative exemplar of its category, by definition, contains better information about its category and thus serves as a better initial prediction (i.e., has high statistical regularity). We ask whether such statistically regular and irregular stimuli elicit differential N300s, as would be hypothesized if this component is indexing hierarchical inference or predictive coding beyond objects.

### Good and bad scenes

We have previously found that good scene exemplars are more readily detected than bad exemplars (Caddigan et al. 2010, 2017); that is, participants are better at discriminating briefly presented and masked intact photographs from fully phase-scrambled versions when those images are good exemplars of their category (i.e., beaches, forests, mountains, city streets, highways, and offices). Good and bad exemplar status was determined with a separate rating task in which participants rated on a 1-5 scale how representative the image was of its category. We took the 60 highest and 60 lowest rated images from each category, and verified that participants were significantly faster and more accurate at categorizing the good scene exemplars than the bad, indicating that our manipulation captured the degree to which the image exemplified the category (Torralbo et al. 2013). Importantly, again, there were no artificially introduced objects in any of the bad exemplars nor were they impoverished or degraded in any way. Instead, their good and bad status derived entirely from how representative they were of the category being depicted. Note that, although category was relevant to the choice of stimuli and whether they were designated good or bad, in Caddigan et al.’ experiments it was completely irrelevant to the intact/scrambled judgement being made (was the stimuli an intact photo or noise?). Nonetheless, participants had significantly higher sensitivity (d’) for good than bad exemplars (Caddigan et al. 2010, 2017), suggesting that with the very brief (34–78 m) masked exposures good exemplars perceptually cohere into a intact photograph sooner than bad exemplars.

Relatedly, the categories of those same good exemplars are better decoded, using fMRI multi-voxel pattern analysis, than are the categories of the bad exemplars in a number of visual areas, including V1 and the parahippocampal place area (PPA; Torralbo et al. 2013). Interestingly, the BOLD signal for those same bad exemplars is larger than that for good exemplars in the PPA (Torralbo et al. 2013), in keeping with predictions from hierarchical predictive coding (i.e., increased activity for the less statistically regular images). The poorer detection with brief presentations, weaker representations in the brain, and greater activity evoked by bad than good scene exemplars make these stimuli good candidates for eliciting a neural signature of hierarchical predictive coding.

### Design of the current experiments

In **Experiment 1**, we recorded scalp EEG while participants viewed good and bad scene exemplars and made a good/bad judgment. If the N300 serves as an index of matching incoming stimuli to learned statistical regularities, then N300 amplitude should be smaller for good exemplars of natural scenes than the bad exemplars. In this first experiment participants viewed the stimuli without any forewarning of what to expect (category and good/bad status were fully randomized; see **Figure 1A**), and all the stimuli were unique images with no repeats in the experiment. If we observe an effect of statistical regularity, then the particular regularity brought online must stem from the current input, as there was no confound of repetition priming or episodic memory.

**Figure 1.**
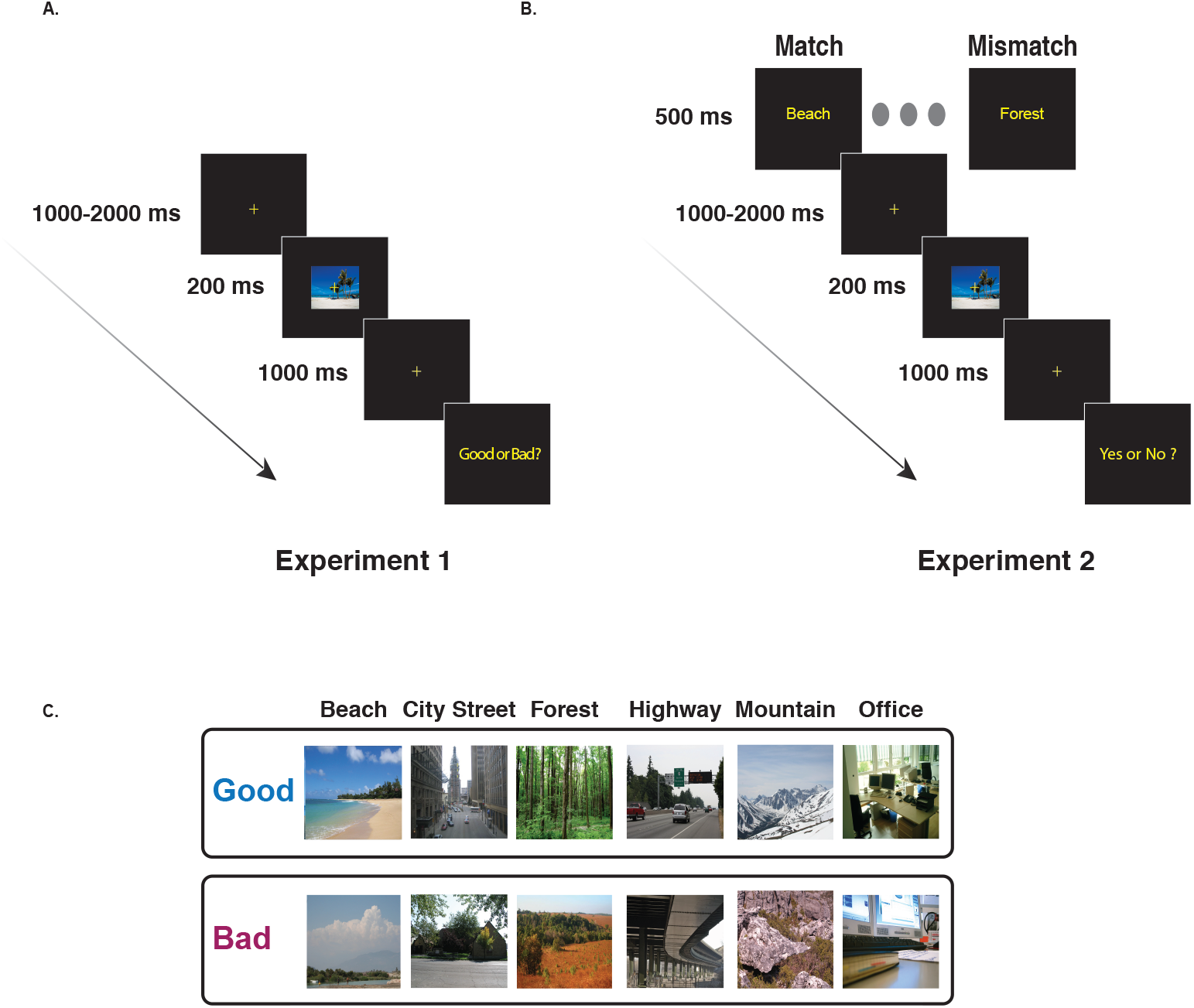
Schematic of one trial in each of the experiments. **A.** In **Experiment 1**, a fixation cross was shown in the center of screen for a randomly chosen interval between 1000-2000 ms. A good or bad exemplar image from one of the six categories was then presented for 200ms, followed by a fixation cross. After a delay of 1000ms, the subjects respond to the question “Good or Bad?” with a button press and the next trial begins. **B.** In **Experiment 2,** the trial sequence is similar to **Experiment 1** with the following differences. At the start of each trial a word cue (e.g., “Beach”) from one of six categories (beaches, city streets, forests, highways, mountains, and offices) is shown. At the end of the trial the subjects make a delayed response, with a button press, to the question “Yes or No?” (“Yes” if the image matches the cue and “No” otherwise) and the next trial begins. Cue validity was kept high (75%) to promote prediction; on 25% of the trials, there is a mismatch between the word cue and the image category. **C.** A sample of good and bad exemplars from each category used in our study.

However, an effective prediction process must also be sensitive to context. Thus, in **Experiment 2** we then manipulated the expectations of the participants at the beginning of each trial by presenting a word cue (e.g., ‘Beach’) that either matched the upcoming scene’s category (on 75% of trials) or mismatched the upcoming image category (e.g., preceding a forest with the ‘Beach’ cue; see **Figure 1B**). If the N300 reflects a PHT process then it should also be sensitive to the particular template (i.e., statistical regularity) activated by the cue. In particular, we would predict that a cue with a 75% validity would activate the statistical regularities associated with the cued category. For images that come from the cued category, then, we should observe smaller N300s for good than bad exemplars, as in Experiment 1, since good exemplars are a better match to the statistical regularities of their category. However, in contrast, when the input image does *not* come from the cued category (i.e., for mismatches), we would predict a reduction or even elimination of the good/bad N300 effect, since neither the good nor bad exemplar would fit well with the cued statistical regularity. For example, good beach exemplars should not systematically provide a better match to the statistical regularities of a forest than a bad beach does. Experiment 2, then, provides a critical test of the idea that the N300 reflects the process of matching input to the currently activated template – i.e., the prediction.

## Materials and Methods

### Participants

The data for **Experiment 1** came from 20 right-handed college-age subjects (mean age = 24.36 years, range = 18 to 33 years, 12 women), and the data for **Experiment 2** from a separate set of 20 right-handed subjects (mean age = 22.44; range 18-30 years; 14 women). In both experiments, participants gave written, informed consent and were compensated for their participation in the study with course credit or cash. The study was approved by the Institutional Review Board of the University of Illinois at Urbana-Champaign. All participants were right-handed, as assessed by the Edinburgh Inventory (Oldfield 1971) and none had a history of neurological disease, psychiatric disorders, or brain damage.

### Materials and Procedures

ERP-eliciting stimuli were pictures of natural scenes from six categories: beaches, forests, mountains, city streets, highways and offices (Figure 1C). In a previous study, these images were collected from the internet and rated for their representativeness of the named category on Amazon Mechanical Turk, with participants answering, e.g., for beaches, ‘‘How representative is this image of a BEACH?’’ for each image, with the interpretation of the term representativeness left to the participants (Torralbo et al. 2013). In a separate experiment, participants were significantly faster and more accurate at categorizing the good exemplars than the bad, further confirming that our manipulation captured the degree to which the image exemplified the category. The 60 top rated images were used as good exemplars for each category, and the 60 lowest rated images were used as bad exemplars for each category (for details on the choice of good and bad exemplars see (Torralbo et al. 2013). Images were resized to 340 × 255 pixels and presented on a black background with a fixation cross at the center. The images were randomly presented at one of three locations: the center of the scene, or with nearest edge 2 degrees to the left or right of fixation, with a total of 120 good images and 120 bad images presented at each location. Here, we report only results for centrally-presented images^1^. The stimuli were all unique images with no repeats in the presentation sequence.

In **Experiment 1**, participants were instructed at the beginning of the study that they would be seeing good and bad exemplars of six scene categories and that their task at the end of each trial was to indicate via button press whether the image was a good or a bad exemplar of its category. Participants first practiced with 9 trials to acclimatize to the task environment, and these images were not repeated in the main experiment. Then, they completed 3 blocks each consisting of an equal number of trials, for a total of 240 centrally presented trials (trials were also presented to the left and right visual fields in each block). The trial counts for centrally presented stimuli, for each category (good and bad combined) are as follows: beaches = 39; cities = 41; forests = 38; highways = 42; mountains = 36; offices = 44.

Participants were seated at a distance of 100 cm from the screen, and the images subtended a visual angle of 7.65° × 5.73° (width × height). Subjects were instructed to maintain fixation on the central fixation cross and to try to minimize saccades and eye blinks during stimulus presentation. As depicted in **Figure 1A**, each trial began with a fixation cross presented on a blank screen for a duration jittered between 1000-2000 seconds (to reduce the impact of slow, anticipatory components on the ERP signal). The scene image, either a good exemplar or a bad exemplar from one of the six categories, was presented for a duration of 200 ms, followed by a fixation cross on a blank screen for 500 ms. At the end of the trial a prompt with “Good or Bad?” was displayed on the screen, and participants pressed one of two response buttons, held in each hand (counterbalanced across participants), to indicate their judgment. The experiment lasted for approximately one hour and fifteen minutes. Subjects were given two five-minute breaks at roughly 25 minutes and 60 minutes from the start of the experiment.

**Experiment 2** was identical to **Experiment 1**, except that each trial began with a word cue, presented for 500 ms (**Figure 1B**), which corresponded to one of the six scene categories used in the experiment: Beach, City Street, Forest, Highway, Mountain, and Office. For each category, we ensured that five trials of each type (good and bad exemplars) were mismatched. There were thus 75% matched trials (15 trials each of good and bad within each of the six scene categories) and 25% mismatched trials, for a total of 180 (90 good, 90 bad) matched trials and 60 mismatched trials (30 good, 30 bad). Overall cue validity was kept high to promote the use of the cue to form expectations about what kind of image would appear next, while still ensuring that we would nevertheless have a sufficient number of mismatch trials to obtain a stable ERP to that condition as well. Instead of making a good or bad judgment, at the end of each trial participants were prompted to respond “yes” or “no,” with a button press, to the question of whether or not they thought that the picture had matched the cue. Hand used to respond “yes” or “no” was counterbalanced.

### ERP Setup and Analysis

EEG was recorded from 26 channels of passive electrodes that were equidistantly arranged on the scalp, referenced online to the left mastoid and re-referenced offline to the average of the left and right mastoids. Additional electrodes placed on the outer cantus of each eye and on the orbital ridge below the left eye were used to monitor saccadic eye movements and blinks. Impedances were kept below 5 KΩ for scalp channels and 10 KΩ for eye channels. The signal was bandpass filtered online (0.02 Hz – 100 Hz) and sampled at 250 Hz. Trials with artifacts due to horizontal eye movements or signal drift were rejected using fixed thresholds calibrated for individual subjects. Trials with blinks were either rejected, or, for subjects with higher numbers of blink artifacts (12 in **Experiment 1** and 8 in **Experiment 2**), were corrected using a blink correction algorithm (Dale 1994). We confirmed that the analytical results were unchanged if blinks were rejected instead of corrected. On average, in **Experiment 1**, 6.83% of good exemplar trials and 9.04% of bad exemplar trials were rejected due to artifacts, and no condition had fewer than 63 trials per subject in the analysis. The average number of retained trials was, for good exemplars, 112 (range 81 to 119) and, for bad exemplars, 109 (range 63 to 120). In Experiment 2, in the match condition, 10.8% of good exemplar trials and 11.09% of bad exemplar trials were rejected due to artifacts and no condition had fewer than 56 trials per subject in the analysis (retained good exemplar trials: mean 80 (63-90); retained bad exemplar trials: mean 80 (56-90)). In the mismatch condition, 10.38% of good exemplar trials and 13.89% of bad exemplar trials were rejected due to artifacts (retained good exemplar trials: mean 27 (19-30); retained bad exemplar trials: mean 26 (19-30)).

ERPs were epoched for a time period spanning 100 ms before stimulus onset to 920 ms after stimulus onset, with the 100 ms prestimulus interval used as the baseline. This processed signal was then averaged for each condition within each subject. A digital bandpass filter (0.2 Hz – 30 Hz) was applied before measurements were taken from the ERPs. Based on prior work showing that the N300 is frontally distributed and occurs between 250 ms to 350 ms (Federmeier and Kutas 2001; Schendan and Kutas 2002, 2003), we measured N300 mean amplitudes in this time window across the 11 frontal electrode sites: MiPf (equivalent to Fpz on the 10-20 system), LLPf, RLPf, LMPf, RMPf, LDFr, RDFr, LMFr, RMFr, LLFr, and RLFr (first letter: R=right, L=left, Mi=midline; second letter: L=lateral, M=medial, D=dorsal; Pf = prefrontal and Fr= frontal); on the 10-20 system, this array spans from Fpz to just anterior of Cz and from mastoid to mastoid laterally, with equidistant coverage. Statistics were computed using R (R Core Team 2020). Specifically, we used the functions t.test, to compute t-tests, and ttestbf (from the package: BayesFactor) to compute Bayes Factors. The t-test and Bayes factor calculations compared the measured condition difference to 0. For within-subject calculations of confidence intervals, we used the function summarySEwithin() that is based on (Morey 2008). The function anovaBF (from the package: BayesFactor) was used to compute Bayes factors for interactions.

For completeness, we also analyzed two ERP components in the time-window after the N300: the N400 and the Late Positive Complex (LPC). Prior work examining the N400 to pictures has shown a frontal distribution (Ganis et al. 1996), and thus we again used the 11 frontal electrode sites, but now in the time-window 350-500 ms. For the LPC we chose posterior sites in the time-window of 500-800 ms based on prior work characterizing the distribution and timing of the LPC (Finnigan et al. 2002).

## Results

### Experiment 1

#### Behavior

To motivate participants to attend to the scenes, we asked participants to make a delayed response on each trial, judging whether the exemplar was a good or bad exemplar of the scene category to which it was presumed to belong. Participants labeled most good exemplars as “good” (mean = 86.2%, std. dev = 13.9%) and labeled bad exemplars as “bad” about half the time (mean = 56.2%, std. dev = 15.6%). All trials (irrespective of the choice of the participants) were used for the planned ERP analyses, but, as described below, we also confirmed that the results hold when conditionalized on subjects’ responses.

#### ERPs

Grand-averaged ERPs at eight representative sites are plotted in **Figure 2**. Responses to good and bad exemplars can be seen to diverge beginning around 250 ms after stimulus onset, with greater negativity for bad exemplars than for good exemplars. The polarity, timing, and frontal scalp distribution of this initial effect is consistent with prior work describing the N300 (Mcpherson & Holcomb, 1999; Schendan & Kutas, 2002, 2003, 2007); see **Supplementary Materials** for a formal distributional analysis.

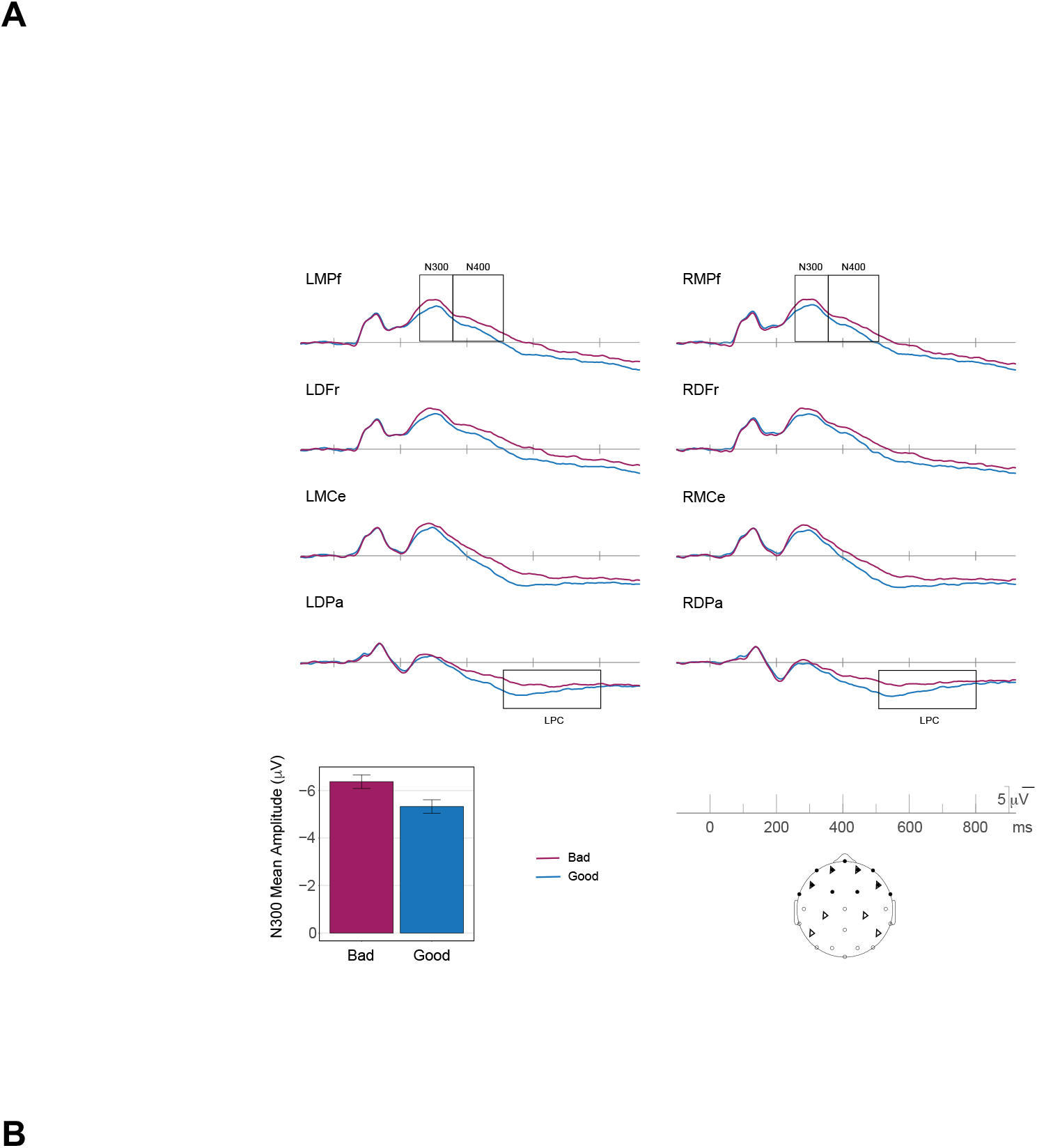

**Figure 2 A.**
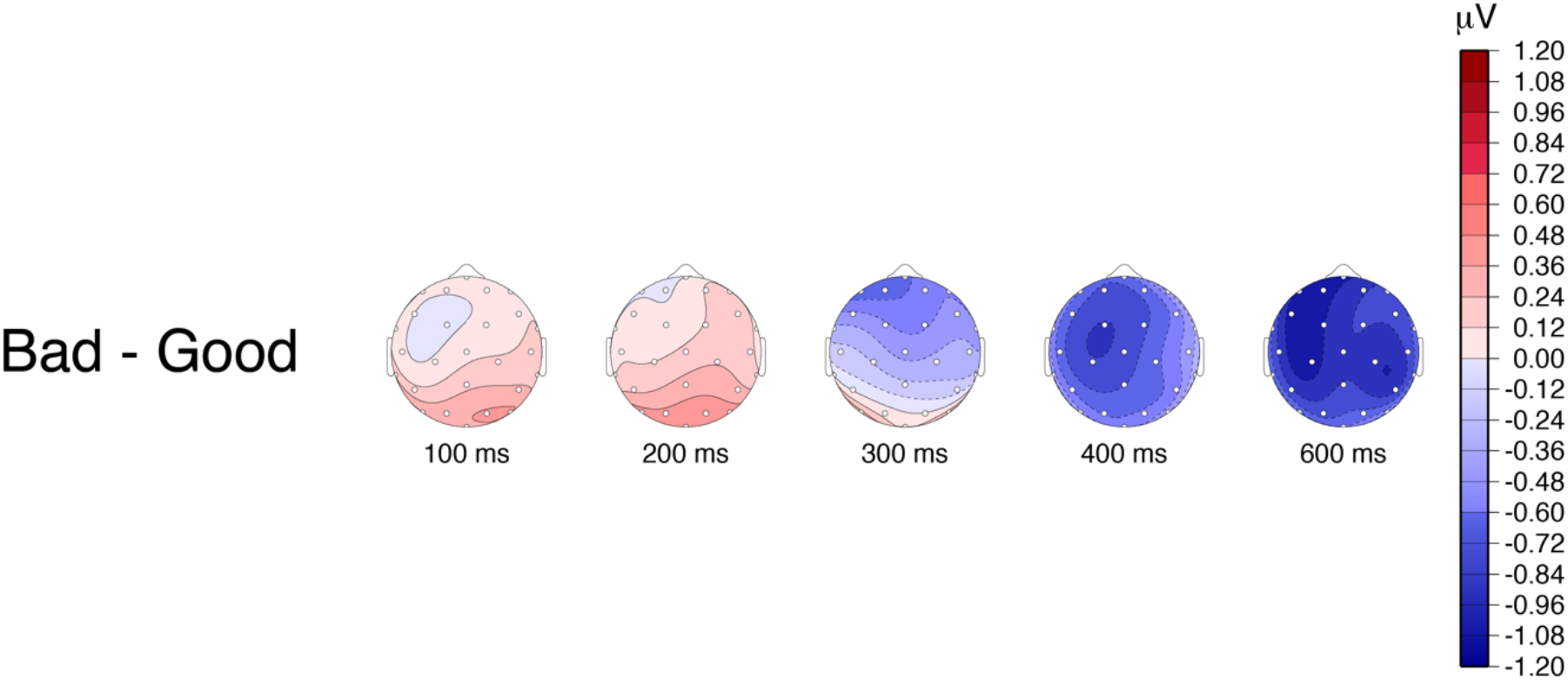
Grand average ERP waveforms for good (blue) and bad (maroon) exemplars in **Experiment 1** are shown at 8 representative electrode sites distributed over the head. Plotted channel locations are marked as triangles on the schematic of the scalp (LMCe and RMCe are just posterior of and lateral to Cz on the 10-20 system). Negative voltage is plotted upwards. The waveforms differ over frontal sites beginning in the N300 time-window (250-350 ms), with greater negativity for bad exemplars as compared to good exemplars. The bar plot gives mean amplitude over the 11 frontal electrode sites (darkened electrode sites on the schematic of the scalp) used for the primary statistical analyses. The error bars plotted are within-subject confidence intervals. N=20. **B.** Topographic plots of the difference waves for the main effect of representativeness (Bad – Good). In the N300 time-window we see a frontal distribution, whereas in the N400 time-window we see a centro-parietal distribution, with a slightly left laterality.

##### N300

To characterize the good/bad effect on the N300, mean amplitudes were measured from all 11 frontal electrode sites between 250 and 350 ms. Bad exemplars elicited significantly larger (more negative) N300 responses (mean = −6.4 μV) than did good exemplars (mean = −5.3 μV); t(19)= −5.4 and Bayes Factor = 747.7 (**Table 1;** for a full distributional analysis see **Supplementary Materials**). In other words, we see the predicted differential response to statistically irregular exemplars (bad exemplars) as compared to the statistically regular exemplars (good exemplars). The larger amplitude for the bad exemplars, as compared to the good exemplars aligns with PHT predictions that would posit greater inference error, and, hence, greater iterative processing for the bad exemplars as compared to the good exemplars. These results also confirm that the N300 indexes a match to statistical regularities of natural scenes and thus extend the validity of the N300 to not only objects, or objects in scene contexts, but more broadly to complex natural scenes.

**Table 1.** **Experiment 1**, mean amplitudes in the N300 time-window (250-350 ms) over 11 frontal electrode sites (see **Figure 2**), along with t-test and Bayes factor values. The N300 response to bad exemplars is more negative (larger) than that to good exemplars. The t-test and Bayes factor calculations compared the within subject Good/Bad difference to 0.

| Condition | N | Mean<br>(μV) | Mean<br>Bad/Good<br>Difference<br>(μV) | Bad/Good<br>Difference 95%<br>C.I. | t(19) | p | Bayes<br>Factor |
| --- | --- | --- | --- | --- | --- | --- | --- |
| Bad | 20 | -6.4±0.61 | -1.05 | -1.46 to -0.64 | -5.4 | 3.3E-05 | 747.7 |
| Good | 20 | -5.3±0.61 | | | | | |

Note: ± values reflect the normed standard deviation within subjects. C.I. = confidence interval.

The above analysis was computed on all trials, to avoid confounding N300 response patterns with the outcome of late stage decision making processes. However, for completeness, we also analyzed the results conditionalized on participants’ responses (i.e., including only good trials judged as good and bad trials judged as bad). This yielded the same effect pattern (Bayes factor for good/bad difference = 5.4; t = −2.89, p = 0.0094). For details see **Supplementary Materials**. We also analyzed the bad exemplar trials, as about half of them were judged as good, and did not see an N300 effect based on participants’ judgements of only the bad exemplars (see **Supplementary Materials)**.

##### Post N300 Components

Although the N300 was the component of primary interest, to more completely characterize the brain’s response to the scenes, we also examined good/bad differences in later time windows encompassing the N400 (350-500 ms) and Late Positive Complex (LPC) (500-800 ms). The details of the analyses and results are provided in the **Supplementary Materials** and summarized here. N400 responses, which index multimodal semantic processing, were larger for bad (−3.3 μV) than for good exemplars (−2.2 μV), suggesting that items that better fit their category allow facilitated semantic access. We note however, that given the similar scalp distribution of the N300 and the N400 to picture stimuli (Ganis et al. 1996), it is difficult to tell where the boundary of the two components might be and thus how much the N400 pattern might be influenced by the preceding N300. LPC responses were larger -- more positive – to good (4.5 μV) than to bad (3.3 μV) exemplars. The LPC amplitude is known to positively correlate with confidence in decision making (Finnigan et al. 2002; Schendan and Maher 2009). Larger LPC responses to good items, therefore, is consistent with the behavioral pattern in which good exemplars were classified more consistently than bad exemplars.

### Experiment 2

As mentioned in the introduction, a predictive coding signal should be sensitive to context. In particular, if the context predicts a specific stimulus category then initial predictions should reflect the statistical regularities associated with the predicted category. The good/bad difference observed in **Experiment 1** was elicited without any expectation regarding the specific category to be presented (i.e., category and good/bad status were completely randomized). Thus, the particular template or statistical regularity with which the image was compared must have been initially elicited by the input itself. This is also the case in almost all previous work examining the N300 to objects. In **Experiment 2**, therefore, we set out to examine whether the N300 is sensitive to expectations induced in the moment by context. We preceded each image with a word cue that either matched or mismatched the upcoming category. If the N300 difference observed in **Experiment 1** reflects the matching of incoming stimuli to learned statistical regularities, we should be able to modulate that difference by activating either the appropriate (match cue) or inappropriate (mismatch cue) statistical regularity. In particular, since neither a good nor a bad exemplar of, e.g., a beach, should be a better match to an inappropriate category (e.g., a forest), we should find that the N300 good/bad difference is reduced or eliminated when the cue mismatches the current category.

#### Behavior

On each trial, participants were asked to respond if the stimulus matched the verbal cue (“Yes” or “No”) via a button press. In the match condition, participants responded “Yes” to good exemplars (mean = 98.7%, std. dev = 2.4%) more often than to bad exemplars (mean= 67.9% and std. dev = 14.6%). In the mismatch condition, wherein the exemplars did not fit the cued category, participants responded “No” to good exemplars (mean = 95.9%, std. dev = 4.6%) more often than to bad exemplars (mean = 94.0% and std. dev = 5.5%). All trials were used for the ERP analyses.

#### ERPs

Scenes elicited an N300 response (**Figure 3**) with similar timing, polarity and scalp distribution to that observed in **Experiment 1**; see the **Supplementary Materials** for a formal distributional analysis. Analyses of N300 mean amplitudes were conducted using the same time window (250-350 ms) and frontal electrode sites as in **Experiment 1**, here comparing good and bad exemplars under the two cueing conditions: match and mismatch.

##### N300

In the match condition, when the scene was congruent with the verbal cue, we replicated the N300 effect of **Experiment 1** for the good and bad exemplars, with a frontally distributed negativity that was larger for the bad exemplars than the good exemplars (**Figure 3, Table 2A, 2B**). Importantly, and as predicted, this N300 difference between good and bad exemplars was notably reduced – indeed, likely absent altogether (Bayes factor 0.31) – in the mismatch condition compared to the match condition (Bayes factor for interaction of Good/Bad x Cuing = 4.0). This is consistent with the idea that the N300 is indexing the fit of the incoming stimulus to the template activated by the verbal cue. That is, neither a good or bad exemplar of category A represents a better match to a template for category B. The same pattern of results is also seen when the analysis is conditioned on subjects’ judgement; i.e., they responded to a cue congruent stimulus as ‘Yes’ and cue incongruent stimulus as “No, see **Supplementary Materials, Table S6**. We note that we chose to discuss the interaction in terms of the good/bad effect being dependent on a matching cue. However, one might also discuss the interaction in terms of the effect of cueing being different as a function of good/bad status. Indeed, the good images show a decrease in the N300 when they are preceded by a match cue than when they are preceded by a mismatched cue (Bayes Factor = 1.2 for good mismatch – good match; t = −1.998, p = 0.06), consistent with the mismatch cue producing a prediction error. In contrast, not only is there little evidence for a cueing effect for bad exemplars (Bayes Factor = 0.68 for bad mismatch – bad match; t = 1.59, p = 0.13) but the difference is numerically in the opposite direction (slightly larger for match).

For completeness, and to compare the N300 in our experiment with its characterization in the existing literature, we also performed an ANOVA across multiple factors: Good/Bad × Cueing (Match/Mismatch) × Anteriority × Laterality × Hemisphere. There was a main effect of Good vs. Bad (bad larger than good; (F(1,19) =15.34) and an interaction between Good/Bad and Cueing (F(1,19) =5.87), with larger Good/Bad effects when the scene matched the cue. The main effect of Cueing was not significant (F(1,19) =0). For details on the distributional analysis see **Supplementary Materials.**

Finally, to ensure that our results are not due to the differential number of trials in the match and mismatch condition, we subsampled the trials in the match condition to be equal to that of the mismatch condition. This subsampling did not change the results **(see Supplementary Materials, Table S7)**.

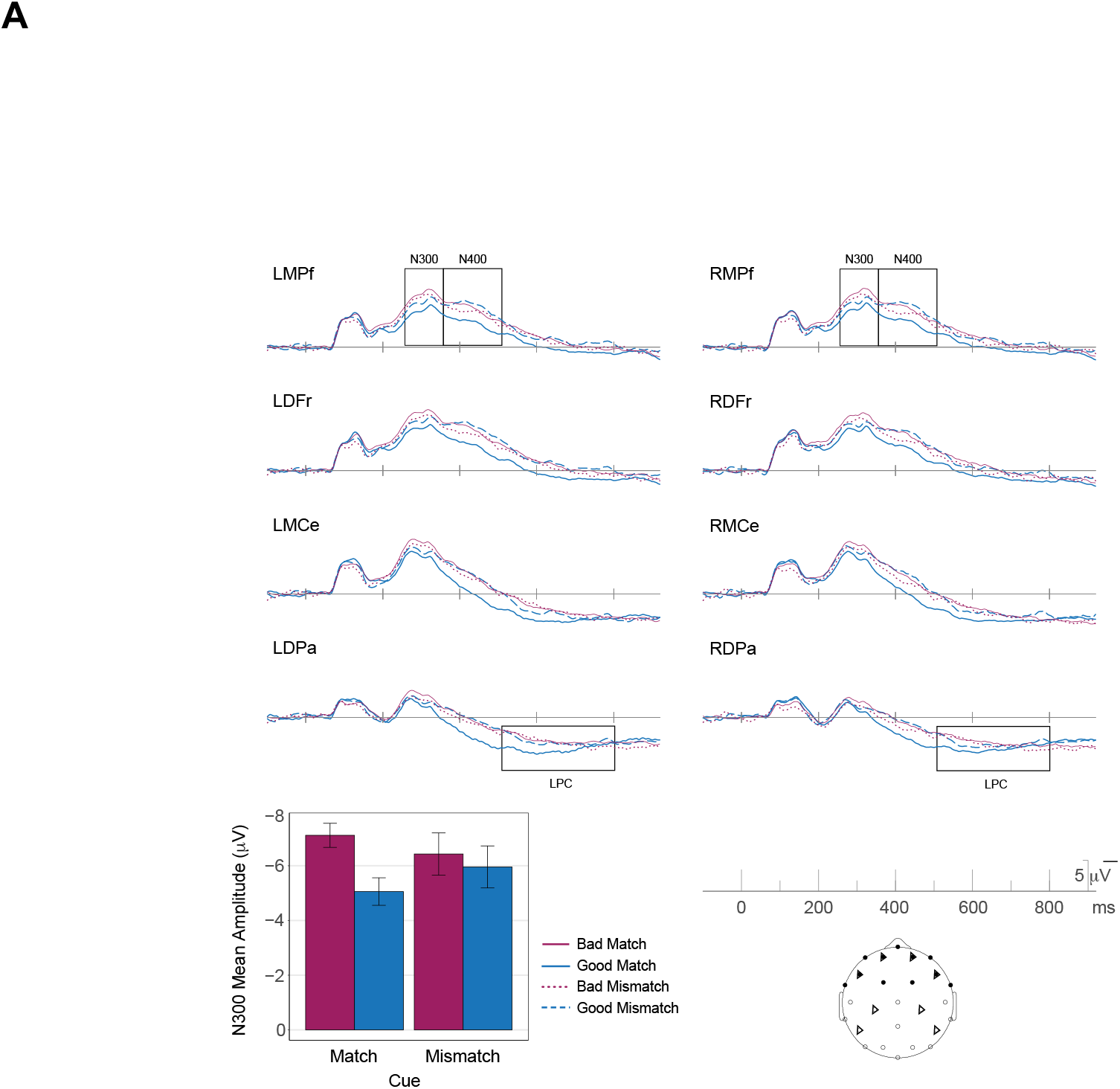

**Figure 3 A.**
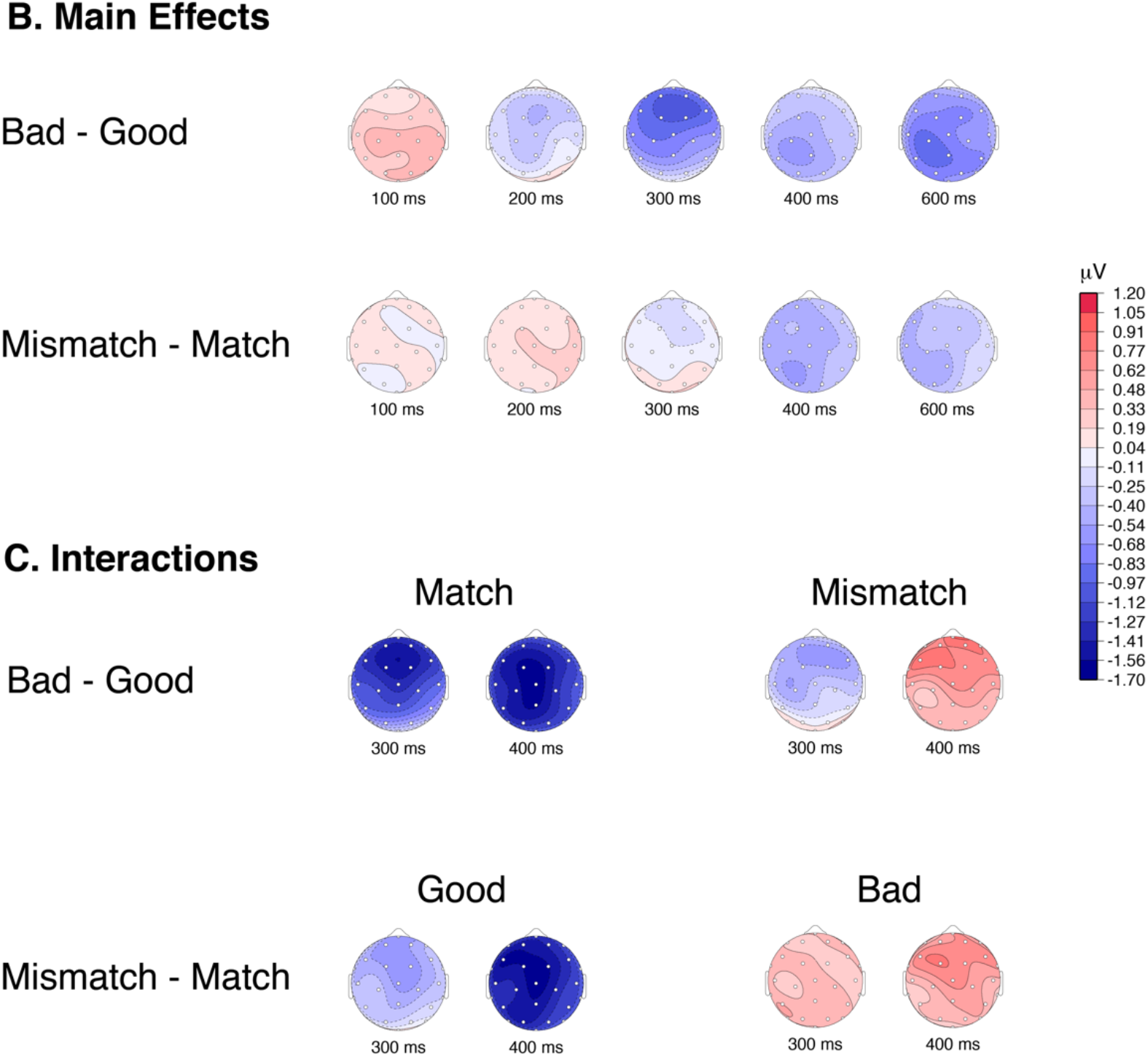
Grand average ERP waveforms for the good-match (solid-blue), bad-match (solid-maroon), good-mismatch (dashed-blue), and bad-mismatch (dotted-maroon) conditions in **Experiment 2** are shown at the same 8 representative electrode sites. In the match condition, responses to good and bad exemplars differ in the N300 time-window (250-350 ms), with greater negativity for bad exemplars as compared to good exemplars, over frontal sites (darkened electrode sites on the schematic of the scalp). In the mismatch condition, the differences between good and bad exemplars on the N300 are diminished/eliminated. The bar plot gives the grand average mean of the ERP amplitude over the 11 frontal electrode sites (darkened electrode sites on the schematic of the scalp) used for the primary statistical analyses (N = 20). The plotted error bars are within-subject confidence intervals. **B.** Topographic plots of the difference waves for the two main effects of representativeness (Bad – Good) and cueing (Mismatch – Match). In the N300 time-window the two main effects are qualitatively similar, with both main effects showing a frontal distribution. The N300 time-window also shows a quantitatively larger effect for the representativeness (Bad – Good) than for the cueing (Mismatch – Match). In the N400 time-window, both effects are centro-parietally distributed with a slight left laterality. **C.** Topographic plots for the difference in the interactions for Good/Bad × Cuing are shown in two interpretations: in terms of the Good/Bad effect – (Bad-Good) × Match and (Bad-Good) × Mismatch; and in terms of the cuing effect -Good × (Mismatch -Match) and Bad × (Mismatch -Match).

**Table 2A.**
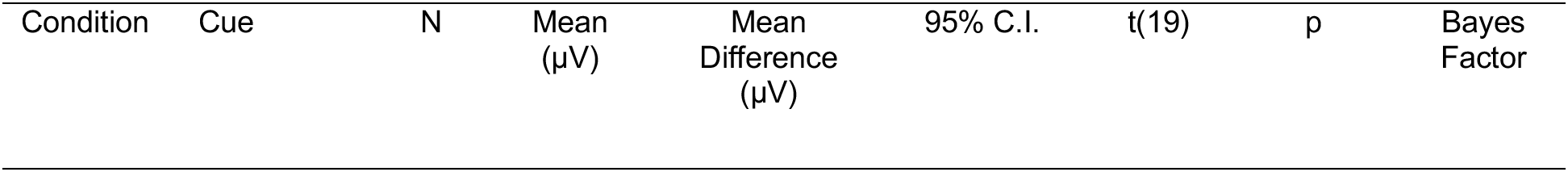
The grand average mean values, in the N300 time-window (250-350 ms), shown for 11 frontal electrode sites (see **Figure 3**), along with t-test and Bayes factor values. There is strong evidence (large Bayes factor) for greater negativity of the N300 for bad exemplars as compared to good exemplars when the cue matches the stimulus. When there is a mismatch between the cue and the stimulus there is no evidence (small Bayes factor) for the difference between good and exemplars in the N300 time-window. The t- test and Bayes factor calculations compared the within subject Good/Bad difference to 0.

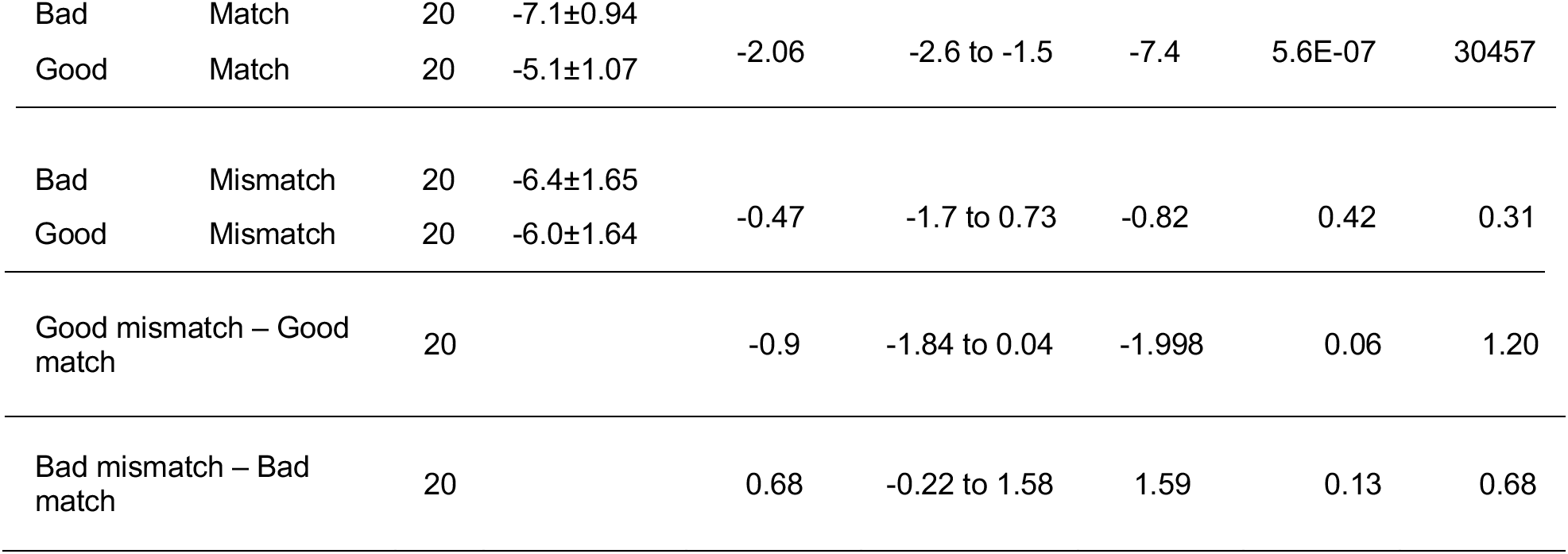

**Table 2B.** The Bayes factor for the main effects and interaction computed using Bayesian ANOVA. This shows that there is evidence for the interaction of Good/Bad x Cueing in **Experiment 2**.

| Effect | Bayes Factor |
| --- | --- |
| Good/Bad | 118.1 |
| Cueing | 0.2 |
| Good/Bad x Cueing | 4.0 |

Note: ± values reflect the normed standard deviation within subjects.

##### Post N300 Components

Again, for completeness, we also examined effects on the N400 (350-500 ms) and Late Positive Complex (LPC) (500-800 ms). These are presented in full in the **Supplementary Materials** and summarized here. Given prior work (reviewed in Kutas and Federmeier 2011), we expected the N400 to be particularly sensitive to the match between the verbal cue and the scene category. Indeed, overall, N400 responses to good scenes that matched the verbal cue were facilitated (more positive: −3.5 μV) than to good scenes that mismatched their cues (−5.6 μV), consistent with the large literature on N400 semantic priming (see **Table S4**). Moreover, we replicated the effect in **Experiment 1**: N400 amplitudes were also larger for bad (−5.3 μV) than for good exemplars (−3.5 μV) in the match condition, although, again, we cannot rule out influence from the prior N300 effects on the observed pattern. We see an interaction of Good/Bad x Cuing in the N400 window (F =13.7; p =0.0015; E =1), with the largest facilitation for good exemplars in the match condition. LPCs were larger (more positive) for good exemplars in the match condition (2.7 μV) compared to both bad exemplars (0.4 μV) in the match condition (replicating **Experiment 1**) and to either scene type in the mismatch condition (Good: 0.2 μV; Bad: 0.9 μV), presumably reflecting the increased ease and confidence of responding to the good match items (see **Table S5**).

## Discussion

In two experiments, we tested whether the N300 component of the ERP has response properties expected for an index of hierarchical predictive coding during late stage visual processing, when global features of the stimulus are being processed. Across many studies, larger (more negative) N300 responses have been observed for conditions that might be characterized as statistically irregular (Pietrowsky et al. 1996; Schendan and Kutas 2002, 2003, 2007; Mudrik et al. 2010; Vo and Wolfe 2013). However, the focus of the literature thus far has been limited to objects, objects in scenes, or artificially degraded stimuli. If the N300 more generally reflects predictive hypothesis testing in later visual processing, then it should be sensitive to statistical regularity outside of the context of object processing and artificial manipulations of global structure. To this end, in Experiment 1 we showed that the N300 is sensitive to the difference between good (statistically regular) and bad (statistically irregular) exemplars of natural scenes. Because none of the scenes we used were degraded, had any misplaced elements, or contained objects that were surprising or violated expectations (e.g., a watermelon instead of the expected basketball; see Mudrik et al. 2010; Vo & Wolfe, 2013), these results strongly link N300 modulations to statistical regularity as such.

Predictive coding posits a larger inference error in processing statistically irregular items (bad exemplars) as compared to statistically regular items (good exemplars), and, consistent with this, N300 responses were larger for the statistically irregular exemplars. Note that the observed pattern cannot be explained by interstimulus perceptual variance (ISPV; Theirry et al., 2007; Schendan and Ganis, 2013). The good exemplars we used have more consistent low-level image statistics, and thus lower ISPV, than the bad exemplars (see Torralbo et al. 2013). Thus, if the pattern were driven by ISPV, we would have expected the good exemplars to elicit larger ERP modulations (see Thierry et al., 2007; Schendan and Ganis, 2013). Instead, we found that the good exemplars have a lower amplitude ERP, consistent with the claim that it is statistical regularity – and not ISPV – that is responsible for the effect.

The data from Experiment 1, in combination with prior experiments, show that the N300 manifests the expected response properties for a general index of predictive coding mechanisms for late stage visual processing (for studies that rule out the N300 indexing early visual processing see Schendan and Kutas 2002; Johnson and Olshausen 2003) of complex objects and scenes. Across the literature, the kinds of stimuli distinguished by the N300 encompass global structure, canonical viewpoints, probable views of objects in scene contexts, and, in our own experiment, the category-level representativeness of the stimuli. We would like to collectively refer to these properties as learned statistical regularities. We mean statistics in the Bayesian sense: The statistical regularities reflect the system’s prior belief. Although frequency of occurrence may be one factor that goes into constructing a regularity, the regularities should be more sophisticated than simple frequency. They should be constructed to maximize the informativeness of the prediction and minimize, on average, the amount of updating needed. Thus, canonicity, prototypicality or representativeness will all be critical determinants of the regularities, as well as frequency or familiarity. A collection of these regularities can be viewed as a template (see also Johnson and Olshausen 2003), constructed to reduce, on average, the prediction error. Thus, we can think of the differences on the N300 component as an indicator of the degree to which an incoming exemplar can be matched with a template, with greater negativity for a stimulus when it doesn't match a template as compared to when it does.

In **Experiment 1**, neither scene category nor exemplar status (good or bad) was predictable from trial to trial, and thus the statistical regularity driving the observed effect must have been acquired over the life time (i.e., learning what does and does not constitute a good exemplar of a category), rather than within the context of the experiment. However, a key attribute of PHT models, of which predictive coding is a popular example, is that the hypotheses that are generated are sensitive to the current context. If the N300 reflects a template matching process, such that the input is compared against a contextually-relevant learned statistical regularity, then the N300 sensitivity to statistical regularity should vary in the moment, as a function of context.

In **Experiment 2,** therefore, we set up expectations for a particular category on each trial using a word cue with high validity, with the aim of pre-activating a particular scene category template. Critically, however, on 25% of trials the scene did not match the cued category. We found that the N300 is indeed sensitive to regularities cued by the current context. When the scenes were congruent with the cued category, we observed a significant effect of statistical regularity (good versus bad) in the N300 time-window, replicating the results from **Experiment 1**. Here the good exemplars provide a better match to the activated template than the bad exemplars, and thus the reduced inference error or iterative matching is reflected in the amplitude of the N300. In the mismatching condition, however, the presented stimulus, whether a good or bad exemplar of its *own* category, does not match the *cued* template (e.g., a “Forest” template has been cued but a good or bad beach scene was presented). In this case, notably, we failed to observe a reliable difference between the N300 to good and bad exemplars. In the language of predictive coding models, similar inference errors would be generated for both statistically regular (good) and irregular (bad) exemplars that mismatch the activated template, as they would both violate the predicted regularities – or, at least, neither good nor bad exemplars of another category should violate the predicted regularities more than the other. Beyond the statistical regularities learned over a lifetime, including our increased familiarity with more prototypical inputs, the N300 shows sensitivity to the specific expectations the visual system has in the moment, generated from the current context.

Others have discussed the use of visual templates in the context of holding information active in memory to afford optimal performance on, e.g., visual matching tasks. In the case of sequential match paradigms, it is assumed that subjects can hold on to a recently seen target object – the “template” in this case – and then use that information to judge subsequent stimuli. Indeed, in these kind of paradigms, differences in anterior ERPs (which may be labeled N2s or N300s; see discussion in Schendan 2019) have been observed between the match and mismatch conditions. Moreover, using a verbal cue for object type (e.g., “dog” followed by an image), Johnson & Olshausen (2003) observed a significant effect of cueing on a frontally-distributed negativity between 150 and 300 ms, which likely is encompassed by what we are calling the N300. Responses were more positive when the image matched the cue compared to when it did not. They did not vary the representativeness of their images, but it is reasonable to assume that they were on average more representative than our bad images, specifically chosen to be less representative. Thus, our results are in accordance with those of Johnson & Olhausen (2003), and extend them, not only to natural scenes, but also by showing that the effect of cuing interacts with sensitivity to statistical regularity. Thus, Experiment 2 brings together two important facets of visual processing on a PHT framework. First, is the fact that the visual system builds templates based on statistical regularities, accumulated over the lifespan, and routinely uses those templates, elicited by the input itself, to guide its iterative processing. Second, then, is that fact that context information (such as a verbal cue) can cause a *particular* template to be activated in advance of the input, biasing processing toward that template.

### The N300 Indexes Perceptual Hypothesis Testing

We can think of visual identification and categorization as a cascade of processes, starting with identification of low level visual features, followed by perceptual grouping of features, and then appreciation of the “whole” visual form of objects and scenes, after which processing moves beyond the visual modality into multi-modal semantics and decision making. PHT mechanisms can work within and across each of these stages. In the context of object processing, prior work on the N300 has posited it as an index of object model selection, an intermediate stage in the process of object identification and categorization (Schendan, 2019; Schendan & Kutas, 2002, 2003, 2007). Having extended the N300 differences to natural scenes, we propose that the N300 reflects PHT mechanisms in this intermediate stage more broadly, not just object selection. Similar to other work (Schendan, 2019), we believe that the N300 reflects processing at the point wherein the input is matched to items in memory with similar perceptual structures. However, our data show that this process is not limited to objects and that it makes use of variety of statistical regularities learned from the world, including those critical for processing both objects and scenes. The broadened view of the N300 as being reflective of a general visual template matching process would suggest that its source be occipitotemporal visual areas. Indeed, the N300 response to objects has been source localized to occipitotemporal visual areas (Schendan & Lucia, 2010; Sehatpour et al., 2006). Although the N300 for scenes has not yet been source localized, a high-density ERP study on scene categorization localized activity in the 200-300 ms time window to these same occipitotemporal visual areas (Greene and Hansen 2020). Similarly, Kaiser and colleagues (2019, 2020), using both fMRI and ERPs, demonstrated a similar sensitivity to intact versus jumbled scenes in the occipital place area and PPA as they did in the N300 time window. Moreover, our prior fMRI work with good and bad scene exemplars (Torralbo et al. 2013) would suggest that the N300 for scenes originates in the PPA, a region known to preferentially process natural scenes (Epstein and Kanwisher 1998). Using the same good and bad scene exemplars as in our experiments, we found that, in the PPA, bad exemplars elicited a greater BOLD signal than good exemplars (Torralbo et al. 2013), mirroring the effect we observed for the N300. Interestingly, in that same PPA region of interest we observed that good exemplars were better decoded than bad exemplars; that is, we were better able to predict the scene category presented on the basis of activity patterns when the scene was a good exemplar than when it was bad in the same region that showed greater activity for the bad exemplar (Torralbo et al. 2013). In other words, it was not the case that reduced activity for good exemplars reflected a weaker representation but instead likely reflected a more efficient representation, an interpretation that aligns nicely with our characterization of the N300 effect as one of visual template matching in occipitotemporal cortex. We suggest that the N300 may be interpreted as a component that reflects the iterative processing, as posited by PHT, in occipitotemporal cortical regions, which helps match previously learned regularities of objects and scenes with the incoming stimulus.

Although we are arguing that the N300 indexes PHT for late stage visual processing of complex visual objects and scenes, it is possible that other components could index PHT at other stages of processing. For example, PHT matching low level sensory features, such as gratings (Kok et al. 2012), to hypotheses about such low level features should occur at earlier stages in the processing hierarchy. Earlier visual sensory components can manifest sensitivity to expected visual features (Boutonnet and Lupyan 2015) or to differences between well-learned visual categories, such as words vs. objects, and faces vs. objects (Schendan et al. 1998) – category comparisons that are thus at a much higher taxonomy than within objects or scenes. Of particular relevance to PHT is the vMMN which, as overviewed in the introduction, temporally precedes the N300 and has been observed in experimental contexts wherein a stream of standard stimuli that share particular low-level visual features (e.g., orientation, color) is occasionally interrupted by the presentation of a target stimulus that carries a featural difference (Stefanics et al. 2014; Oxner et al. 2019). Thus, the vMMN is sensitive to the context of recent exposure to low-level visual information, possibly reflecting PHT processes at that lower level.

The N300, instead, does not modulate with low-level differences and manifests sensitivity to both regularities established through long-term experience and knowledge-based expectations derived from semantic contextual information. It may thus index a late stage of visual PHT, at the transition into multimodal, semantic processing. Immediately after the N300, ERP responses to complex objects and scenes are characterized by an N400, which we also observe in our experiment. The N400 is widely accepted as a signature of multi-modal semantic processing, elicited by not only visual words and pictures, but also meaningful stimuli in other modalities (see review Kutas & Federmeier, 2011), whereas the N300 seems to be about visual perceptual structure (Schendan, 2019; Schendan & Kutas, 2002, 2003, 2007). In some cases, it may be difficult to disentangle the precise contributions of the N300 and N400 to observed effects of object categorization and match to object knowledge (Gratton et al., 2009; Schendan, 2019; Schendan & Maher, 2009) since the N400 is known to be sensitive to the fit between, e.g., a picture and its context (Ganis et al. 1996; Federmeier and Kutas 2002). Importantly, however, this does not impact the critical effect of our good versus bad scenes, as neither contain contextually inappropriate items, nor, in **Experiment 1,** did we set up any context prior to an image (i.e., the scene category is unpredictable).

## Conclusion

In a set of experiments we have provided support for the hypothesis that the N300 component is an index of PHT at the level of whole-objects and scenes. Using statistically regular and irregular exemplars of natural scenes, we showed that items that are a poorer match to our learned regularities for types of scenes – and, thus, inputs that should lead to larger inference errors in a predictive coding framework – indeed evoked a larger N300 amplitude compared to statistically regular exemplars, even when the upcoming scene category was not predictable. We further showed, not only that N300 responses to scenes are modulated by context – such as the scene category predicted by a verbal cue -- but that they behave as expected for a template matching process in which statistically regular images procure their advantage by virtue of matching the current contextual prediction.

Our work thus not only extends prior work on the N300 to natural scenes but it suggests that the N300 reflects a general template/model selection process of the sort proposed by PHT models, such as predictive coding. We propose that the N300 indexes visual inference processing in a late visual time-window that occurs at the boundary between vision and the next stage of multi-modal semantic processing. Further studies will be needed to explore the full range of the N300 response. For example, does it require that the object or scene is attended or might it proceed more automatically? Can it be modulated by contexts set up in different modalities (e.g., auditory inputs: speech, sounds)? Regardless, we propose that the N300 can serve as a useful marker of knowledge guided visual processing of objects and scenes, with templates based on prior knowledge serving as hypotheses for visual inference as posited by PHT.

## Acknowledgments

We would like to thank Yanqi Zhang for assistance with running subjects in **Experiment 1**, and Resh Gupta, and Nirupama Mehrotra for helping with **Experiment 1** data collection. We also thank Rami Alsaqri, Johan Saelens, Daria Niescierowicz, and Benjamin D. Schmitt for helping with data collection in **Experiment 2**.

## Footnotes

1 The laterally presented scenes were included to separately answer questions about hemispheric biases in scene processing that are outside the scope of this manuscript. Because ERP waveforms for laterally presented stimuli have important morphological differences compared to those from centrally presented stimuli, the data from the two presentation conditions cannot be combined.

