## Supplementary Materials for "The N300: An Index For Predictive Coding Of Complex Visual Objects and Scenes"

### Experiment 1

#### N300 Distributional Analysis

To compare the N300 in our experiment with its characterization in the existing literature, we performed an ANOVA in the time-window 250-350 ms with the factor of representativeness (Good/Bad) and dividing the 16 scalp channels into 3 additional factors: 2 levels of Hemisphere (right and left scalp sites), 2 levels of Laterality (lateral and medial scalp sites), and 4 levels of Anteriority (prefrontal, frontal, central/parietal, and occipital scalp sites). In addition to confirming the main effect of Good vs. Bad (bad larger than good) ($F(1,19) = 11.97$; $p=0.0026$, $E=1$), there were interactions of Good/Bad with both Laterality ($F(1,19)=7.35$; $p=0.014$; $E=1$) and Anteriority ($F(3,57) = 27.86$; $p<0.0001$; $E=0.5336$), as well as three-way interactions: Good/Bad x Hemisphere x Anteriority ($F(3,57)=3.13$; $p=0.0324$; $E=0.8353$), and Good/Bad x Laterality x Anteriority ($F(3,57)=6.79$; $p=0.0005$; $E=0.7571$). Overall, N300 responses were observed over both left and right hemisphere sites and were largest over the front of the head and larger over medial compared to lateral electrode sites. Over frontal sites, the effect of Laterality was more pronounced and there was a tendency for larger effects over left compared to right hemisphere sites. These results show that the topographic distribution of the N300 effect for natural scenes has a similar distributional profile as the N300 for objects (Schendan & Kutas, 2002, 2003, 2007).

#### Post N300 Components

Analyses of the N400 and LPC are presented in Tables S1 and S2.

**Table S1.** The grand average mean values in the N400 time-window (350-500 ms), shown for 11 frontal electrode sites along with t-test and Bayes factor values. The N400 for the bad exemplars is larger (more negative) than that for the good exemplars. The t- test and Bayes factor calculations compared the within-subject Good/Bad difference to 0.

| Condition | N | Mean (μV) | Bad/Good Difference Mean (μV) | Bad/Good Difference 95% C.I. | t(19) | p | Bayes Factor |
|---|---|---|---|---|---|---|---|
| Bad | 20 | -3.3±0.96 | -1.14 | -1.78 to -0.50 | -3.74 | 0.0014 | 27.3 |
| Good | 20 | -2.2±0.96 | | | | | |

Note: ± values reflect the normed standard deviation within subjects.

**Table S2:** The grand average mean values, in the LPC time-window (500-800 ms), shown for 15 posterior electrode sites (LMCe, RMCe, LDCe, RDCe, MiCe, MiPa, LLTe, RLTe, LDPa, RDPa, LLOc, RLOc, LMOc, RMOc, MiOc) along with t-test and Bayes factor values. The LPC for bad exemplars has a smaller mean amplitude than for the good exemplars. The t- test and Bayes factor calculations compared the within subject Good/Bad difference to 0.

| Condition | N | Mean (μV) | Bad/Good Difference Mean (μV) | Bad/Good Difference 95% C.I. | t(19) | p | Bayes Factor |
|---|---|---|---|---|---|---|---|
| Bad | 20 | 3.3±1.1 | -1.17 | -1.91 to -0.43 | -3.3 | 0.0036 | 12.1 |
| Good | 20 | 4.5±1.1 | | | | | |

Note: ± values reflect the normed standard deviation within subjects.

### ERP Analysis Conditioned on Participants' Judgements

To examine the N300 effect conditionalized on participants' explicit judgements, we repeated the Bayes factor analysis, but now only using trials wherein participants' judgments aligned with the condition designation (i.e., good exemplars judged as good and bad exemplars judged as bad). As can be seen in **Table S3**, using the same time window and same set of electrode sites, we again find a good/bad N300 effect for this subset of trials (Bayes Factor = 5.4; t = -2.89, p = 0.009). The post-N300 components, the N400 and LPC, also show significant effects for this subset of trials, albeit with reduced Bayes factors.

**Table S3.** The grand average mean values along with t-test and Bayes factor values, only for trials in which participants marked good exemplars as good or bad exemplars as bad, for the N300 and post N300 components. This analysis was carried out at identical electrode sites to the corresponding analyses in Tables 1, S1, and S2. The t- test and Bayes factor calculations compared the within-subject Good/Bad difference to 0.

| ERP | Condition | N | Mean (μV) | Bad/Good Difference Mean (μV) | Bad/Good Difference 95% C.I. | t(19) | p | Bayes Factor |
|---|---|---|---|---|---|---|---|---|
| N300 | Bad | 20 | -6.4±1.03 | -0.95 | -1.63 to -0.26 | -2.89 | 0.0094 | 5.4 |
| | Good | 20 | -5.4±1.03 | | | | | |
| N400 | Bad | 20 | -3.2±1.51 | -1.02 | -2.01 to -0.02 | -2.13 | 0.046 | 1.5 |
| | Good | 20 | -2.1±1.51 | | | | | |

| | | | | | | | | |
|---|---|---|---|---|---|---|---|---|
| LPC | Bad | 20 | 3.2±1.67 | -1.20 | -2.3 to -0.09 | -2.26 | 0.0359 | 1.8 |
| | Good | 20 | 4.4±1.67 | | | | | |

**Comparing the N300 to Bad Exemplars Judged as Good or Bad**

Bad exemplars (designated based on large-scale rating data) were explicitly judged to be bad by participants in this study about half the time (mean = 56.2%, std. dev = 15.6%). We compared the N300 amplitude to these exemplars based on participants' judgments and found no evidence that the N300 differs for bad exemplar trials that subjects responded to as "bad" (-6.35 μV) vs. "good" (-6.37 μV) (Bayes Factor = 0.23; t(19) = 0.04; p= 0.97). Participant judgements also did not reliably modulate either the N400 (Bayes Factor = 0.23; t(19) = 0.16, p = 0.88) or the LPC (Bayes Factor = 0.29; t(19) = -0.71; p = 0.48). Thus, we see no indication that the ERP patterns were importantly affected by participants' explicit judgments, although we note that this analysis is conducted on half the trials of the main analysis. Because judgments for the good exemplars were much more consistent (mean judged to be "good" = 86.2%, std. dev = 13.9%), there were insufficient trials to examine the impact of participant judgment for these items.

**Experiment 2**

**N300 Distributional Analysis with Cuing**

A novel feature of **Experiment 2** was the use of a verbal precue. To see how the distributional properties of the N300 effect under cued conditions compare to that in prior work, we performed an ANOVA of the Cueing (match/mismatch) and Good/Bad factors using the same electrode sites as in the ANOVA analysis for **Experiment 1**, adding the identical electrode factors: 2

levels of Hemisphere (right and left scalp sites), 2 levels of Laterality (lateral and medial scalp sites), and 4 levels of Anteriority (prefrontal, frontal, central/parietal, and occipital scalp sites). We replicate the main effect of Good and Bad from **Experiment 1,** with larger N300 responses to bad exemplars than to good; $F(1,19) = 15.34$; $p= 0.0009$, E=1. The topographic distribution of this effect was also similar to that in **Experiment 1** and in the larger N300 literature: N300 effects were observed over both left and right hemisphere sites, with the left hemisphere showing somewhat larger effects as compared to the right hemisphere (Good/Bad x Hemisphere ($F(1,19) =5$; $p= 0.0375$, E=1) and were larger over medial than lateral sites (Good/Bad x Laterality ($F(1,19) =21.1$; $p =0.0002$, E=1) and larger over the front of the head (Good/Bad x Anteriority ($F(3,57) =7.13$; $p =0.0004$, E=0.4030).

We also found an interaction of Good/Bad x Cueing ($F(1,19) =5.87$; $p= 0.0255$, E=1), with large N300 effects when the stimuli matched the cue, but negligible effects in the mismatch condition. There was a 3-way interaction of Good/Bad x Cueing x Laterality ($F(1,19) =4.83$; $p= 0.0406$, E=1), because the tendency for N300 effects to be larger over medial than lateral sites was more apparent in the match condition, which showed strong N300 effects.

**N400: The Effects of Cuing**

The N400 is known to be affected by semantic expectancy. To confirm that pattern and examine its interaction with Good/Bad status in **Experiment 2**, we performed an ANOVA analysis in the N400 time-window (350-500 ms). Consistent with the larger literature, we found a main effect of Cuing ($F =5.43$; $p =0.03$; E =1) with a smaller N400 amplitude to stimuli that matched the cue compared to the those that mismatched. We also found a significant interaction of Good/Bad x Cuing ($F =13.7$; $p =0.0015$; E =1), with the good exemplars in the match condition having the smallest amplitude as compared to the good and bad exemplars in the match- and- mismatched condition. Note that we do not claim that these effects in the N400 time window are completely independent of the preceding N300 effects in our experiment. Prior work has shown that the N300 and N400 are functionally dissociable (Federmeier & Kutas, 2002; Gratton et al., 2009), but their similar response pattern to, e.g., incongruent and congruent items, can make separating them challenging under some circumstances (Draschkow et al., 2018). We also computed the Bayes factor for the N400 in the match and mismatch conditions (**Table S4**) and we see strong evidence for the N400 in the match condition (Bayes factor 1287.4) as compared to the mismatch condition (Bayes factor 0.56).

**Table S4.** The grand average mean values, in the N400 time-window (350-500 ms), shown for 11 frontal electrode sites. The t- test and Bayes factor calculations compared the within subject Good/Bad difference to 0.

| Condition | Cue | N | Mean (μV) | Difference (μV) | Bad/Good Difference 95% C.I. | t(19) | p | Bayes Factor |
|---|---|---|---|---|---|---|---|---|
| Bad | Match | 20 | -5.3±0.98 | -1.82 | -2.5 to -1.15 | -5.67 | 1.8E-05 | 1287.4 |
| Good | Match | 20 | -3.5±1.41 | | | | | |
| Bad | Mismatch | 20 | -4.7±1.65 | 0.91 | -0.42 to 2.25 | -1.44 | 0.17 | 0.56 |
| Good | Mismatch | 20 | -5.6±2.06 | | | | | |

Note: ± values reflect the normed standard deviation within subjects.

**LPC**

For the LPC, we examined differences in the 500-800 ms time window, encompassing the Late Positive Complex (LPC), over 15 posterior sites (identical to the sites for the analysis in **Experiment 1**). We replicate the main effect of Good/Bad with a larger LPC amplitude for good as compared to bad exemplars (F = 4.84; p = 0.0403; E =1). The LPC is known to index confidence in decision making (Finnigan et al., 2002) and the main effect of Cuing in **Experiment 2** aligns with this understanding**,** with the match condition showing a larger LPC amplitude as compared to the mismatch condition (F = 19.69; p =0.0003; E=1). The interaction of Good/Bad x Cuing is also significant (F =19.82; p =0.0003; E =1) with the good match having the largest LPC amplitude as compared to the Good mismatch, Bad match, and Bad mismatch conditions. We also computed the Bayes factor for the LPC in the match and mismatch conditions (Table S5) and we see strong evidence for the LPC in the match condition (Bayes factor 9014.4) as compared to the mismatch condition (Bayes factor 0.41).

**Table S5.** The grand average mean values, in the LPC time-window (500-800 ms), shown for 15 posterior electrode sites. The t- test and Bayes factor calculations compared the Good/Bad difference to 0.

| Condition | Cue | N | Mean (μV) | Mean Bad/GoodDifference (μV) | Bad/Good Difference 95% C.I. | t(19) | p | Bayes Factor |
|---|---|---|---|---|---|---|---|---|
| Bad | Match | 20 | 0.4±0.98 | -2.24 | -2.94 to -1.54 | -6.69 | 2.1E-06 | 9014.4 |
| Good | Match | 20 | 2.7±1.09 | | | | | |
| Bad | Mismatch | 20 | 0.9±1.66 | 0.68 | -0.56 to 1.93 | 1.15 | 0.26 | 0.41 |
| Good | Mismatch | 20 | 0.2±1.61 | | | | | |

Note: ± values reflect the normed standard deviation within subjects.

**ERP Analyses Conditioned on Participants' Judgements**

As for Experiment 1, we also examined Good/Bad effects conditionalized on participants' explicit judgements, only including exemplars on which subjects judgement was congruent with the category cue; i.e., they responded to a cue congruent stimulus as 'Yes' and cue incongruent stimulus as "No", for both good and bad exemplars. As can be seen in **Table S6**, we again found that explicit judgments did not notably impact the ERP patterns.

**Table S6.** The grand average mean values along with t-test and Bayes factor values, only for trials in which participants responded to a cue congruent stimulus as 'Yes' and cue incongruent stimulus as "No", for both good and bad exemplars. This analysis was carried out at identical electrode sites to the corresponding analyses in Tables 2, S1, and S2. The t- test and Bayes factor calculations compared the within-subject Good/Bad difference to 0.

| ERP | Condition | Cue | N | Mean (μV) | Difference (μV) | Bad/Good Difference 95% C.I. | t(19) | p | Bayes Factor |
|---|---|---|---|---|---|---|---|---|---|
| N300 | Bad | Match | 20 | -7.2±1.18 | -2.15 | -2.98 to -1.33 | -5.45 | 2.9E-05 | 835.8 |
| | Good | Match | 20 | -5.1±1.24 | | | | | |
| | Bad | Mismatch | 20 | -6.5±1.68 | -0.55 | -1.72 to 0.63 | -0.98 | 0.34 | 0.35 |
| | Good | Mismatch | 20 | -5.9±1.48 | | | | | |
| N400 | Bad | Match | 20 | -5.3±1.18 | -1.71 | -2.6 to -0.82 | -4.02 | 0.0007 | 48.2 |
| | Good | Match | 20 | -3.6±1.50 | | | | | |

| | | | | | | | | | |
|---|---|---|---|---|---|---|---|---|---|
| | Bad | Mismatch | 20 | -4.7±1.62 | 0.92 | -0.37 to 2.21 | 1.49 | 0.15 | 0.6 |
| | Good | Mismatch | 20 | -5.6±1.91 | | | | | |
| LPC | Bad | Match | 20 | 0.6±1.22 | -2.05 | -2.85 to -1.25 | -5.35 | 3.6E-05 | 689.4 |
| | Good | Match | 20 | 2.6±1.13 | | | | | |
| | Bad | Mismatch | 20 | 1.0±1.76 | 0.79 | -0.42 to 2.01 | 1.36 | 0.19 | 0.5 |
| | Good | Mismatch | 20 | 0.2±1.49 | | | | | |

Note: ± values reflect the normed standard deviation within subjects.

**N300: Subsampling the Match Trials**

The mismatch condition in **Experiment 2** had fewer trials and therefore possibly a lower signal-to-noise-ratio as compared to the match condition. To verify that we see evidence for the Good/Bad effect in the match condition even when trial numbers are equated to the mismatch condition, we randomly subsampled, in the match condition, 30 trials from the good exemplars and 30 trials from the bad exemplars and recomputed the statistics at identical frontal electrode sites. Even with just this subset of trials, we find that N300 amplitudes are larger for bad than for good exemplars in the match condition (Bayes Factor = 35.6; $t=-3.9$; $p=0.001$; **Table S7**). Moreover, when we combine the sampled good data with the mismatch data, there is still an interaction between Cuing and the Good/Bad effect (Bayes Factor = 3.1), such that the Good/Bad effect is reduced under mismatch compared to match conditions.

**Table S7.** The grand average mean values for the N300 and post N300 components for subsampled good and bad exemplars (30 trials each) in the match condition, computed at identical electrode sites as those used in **Table 2**, **S4** and **S5**. The t- test and Bayes factor calculations compared the within subject Good/Bad difference to 0.

| ERP | Condition | Cue | N | Mean (μV) | Difference (μV) | Bad/Good Difference 95% C.I. | t(19) | p | Bayes Factor |
|---|---|---|---|---|---|---|---|---|---|
| N300 | Bad | Match | 20 | -7.9±1.84 | -2.42 | -3.74 to -1.1 | -3.9 | 0.001 | 35.6 |
| | Good | Match | 20 | -5.5±1.77 | | | | | |
| N400 | Bad | Match | 20 | -5.8±1.53 | -1.82 | -3.03 to -0.62 | -3.16 | 0.0051 | 8.99 |
| | Good | Match | 20 | -4.0±1.89 | | | | | |
| LPC | Bad | Match | 20 | 0.3±1.79 | -1.99 | -3.18 to -0.81 | -3.52 | 0.002 | 18.0 |
| | Good | Match | 20 | 2.3±1.61 | | | | | |

Note: ± values reflect the normed standard deviation within subjects.

**Supplementary Materials: References**

Draschkow, D., Heikel, E., Võ, M. L.-H., Fiebach, C. J., & Sassenhagen, J. (2018). No evidence from MVPA for different processes underlying the N300 and N400 incongruity effects in object-scene processing. *Neuropsychologia*, *120*, 9–17. https://doi.org/10.1016/j.neuropsychologia.2018.09.016

Federmeier, K. D., & Kutas, M. (2002). Picture the difference: Electrophysiological investigations of picture processing in the two cerebral hemispheres. *Neuropsychologia*, *40*(7), 730–747. http://kutaslab.ucsd.edu/people/kutas/pdfs/2002.N.730.pdf

Finnigan, S., Humphreys, M. S., Dennis, S., & Geffen, G. (2002). ERP 'old/new'effects: Memory strength and decisional factor (s). *Neuropsychologia*, *40*(13), 2288–2304. http://www.sciencedirect.com/science/article/pii/S0028393202001136

Gratton, C., Evans, K. M., & Federmeier, K. D. (2009). See what I mean? An ERP study of the effect of background knowledge on novel object processing. *Memory & Cognition*, *37*(3), 277–291. https://doi.org/10.3758/MC.37.3.277

Schendan, H. E., & Kutas, M. (2002). Neurophysiological evidence for two processing times for visual object identification. *Neuropsychologia*, *40*(7), 931–945. http://www.sciencedirect.com/science/article/pii/S0028393201001762

Schendan, H. E., & Kutas, M. (2003). Time course of processes and representations supporting visual object identification and memory. *Journal of Cognitive Neuroscience*, *15*(1), 111–135. http://www.mitpressjournals.org/doi/abs/10.1162/089892903321107864

Schendan, H. E., & Kutas, M. (2007). Neurophysiological evidence for the time course of activation of global shape, part, and local contour representations during visual object categorization and memory. *Journal of Cognitive Neuroscience*, *19*(5), 734–749. http://www.mitpressjournals.org/doi/abs/10.1162/jocn.2007.19.5.734
